# Mechanophenotype of synovial fibroblasts correlates with patients’ pathotypes and treatment response in Rheumatoid Arthritis

**DOI:** 10.64898/2026.10.01.755901

**Authors:** Florencia Diaz, Teresa Debatin, Lena Messner, Célia Mouchel dit Grainville, Jorge Andrés, Florian Riethmüller, Ria Vaishampayan Vikrant, Raffaele Fronza, Fereydoon Taheri, Zhe Wang, Christine Selhuber-Unkel, Federico Colombo

**Affiliations:** Institute for Molecular Systems Engineering and Advanced Materials (IMSEAM), Ruprecht-Karls-Universität Heidelberg, Im Neuenheimer Feld 225, 69120 Heidelberg, Germany; AI and Computational Biomedicine Unit, Department of Medical Sciences, University of Torino, Italy

**Keywords:** Rheumatoid arthritis, Mechanical phenotype, 3D cell culture, Treatment response, Multiomics

## Abstract

The mechanical features of synovial fibroblasts may encode clinically relevant information for the management of rheumatoid arthritis. Indeed, it remains poorly understood the links between cellular and microenvironment mechanics with patient stratification and therapeutic response. Here we show that combining organotypic synovial cultures with multimodal mechanophenotyping and transcriptomics identifies distinct ‘soft’ and ‘stiff’ fibroblast states. Healthy and rheumatoid-arthritis fibroblasts were cultured in two or three dimensions (organotypic cell cultures), alone or with endothelial cells, and characterized by high-throughput nanoindentation, picobalance measurements, digital holographic microscopy, Brillouin microscopy and RNA sequencing. Three-dimensional culture reduced single-cell Young’s modulus and total mass while increasing dry-mass surface density; endothelial co-culture increased fibroblast stiffness and the longitudinal modulus of intact organotypic cultures. Stiffness-based grouping revealed transcriptional programmes related to extracellular-matrix remodelling, cytoskeletal contractility and TGF-β, WNT and Hippo signalling. Mapping these signatures to publicly available rheumatoid-arthritis clinical transcriptomic datasets associated the ‘stiff’ state with fibrotic and myeloid-fibrotic pathotypes and treatment-refractory programmes, whereas the ‘soft’ state aligned with lymphoid pathotypes and rituximab- or tocilizumab-responsive programmes. These findings support mechanophenotype-associated transcriptional signatures as potential biomarkers for rheumatoid arthritis stratification.

## Introduction

The mechanical properties of cells and their interactions with the extracellular matrix (ECM) are increasingly recognized as central regulators of tissue homeostasis, immune responses, and disease pathogenesis. At the core of this interplay is mechanotransduction, a process by which cells convert mechanical stimuli into biochemical signals, modulating key cellular behaviors such as migration, differentiation, and proliferation^1–7^. These cues are particularly significant in chronic inflammatory diseases like rheumatoid arthritis (RA) and Osteoarthritis (OA), where both ECM mechanics and cellular biophysical properties are profoundly altered^8–11^. In healthy tissues, ECM stiffness and architecture are tightly regulated to maintain physiological function, yet in RA, synovial tissues undergo extensive remodeling. This is marked by aberrant ECM stiffening, persistent inflammation, and the emergence of hyperplastic and invasive fibroblast-like synoviocytes (FLS)^12–15^. These RA-FLS acquire a transformed tumor-like phenotype characterized by increased migratory and invasive capacity, cytoskeletal remodeling, and marked resistance to apoptosis^16–18^. In general, mechanosensitive pathways, including Hippo/YAP and Notch, are well known to be implicated in the regulation of cell response to altered ECM mechanics and intercellular interactions^19–22^. Inflammatory cues, including tumor necrosis factor and toll-like receptors, trigger a variety of pathways that promote macrophage, fibroblast, and endothelial cell interactions that sustain inflammation over time and eventually joint damage^23^. Traditional two-dimensional (2D) in-vitro cultures, while widely used, fail to capture the complexity of these mechanobiological dynamics and pathways, often omitting the essential 3D spatial context in which cells experience force and structure. In contrast, three-dimensional (3D) culture models offer physiologically relevant platforms for mimicking the in-vivo microenvironment, enabling more accurate insights into mechanotransduction and cell–ECM interactions^24–26^. We have previously demonstrated that varying the frequency of mechanical stress in 3D cell cultures significantly alters actin organization in human synovial fibroblasts, highlighting the importance of biomechanical context in shaping the function of these cells^27^. RA pathogenesis is further compounded by endothelial cell (EC) involvement, which contributes to synovial vascularization and supports fibroblast expansion through paracrine and juxtacrine signaling. Recent studies have shown that Notch signaling governs this crosstalk, controlling fibroblast identity and spatial patterning within the joint^28–30^. Despite these advances, the correlation of the mechanical phenotype of synovial fibroblasts with molecular signatures, patient stratification and therapeutic outcomes remains poorly explored and considered.

To address this, we used a 3D organotypic co-culture model incorporating human fibroblast-like synovial cells from either a healthy donor (HFLS) and from an RA patient (RA-HFLS), alongside human umbilical vein endothelial cells (HUVECs), to recapitulate the biomechanical and cellular complexity of the inflamed synovium. This platform allows us to investigate how the mechanical forces from the ECM 3D environment, and fibroblast–endothelial interactions converge to shape FLS biomechanics in RA. Using this system, we identified two distinct biomechanical phenotypes “soft” and “stiff” among HFLS, each associated with a specific genetic signature. These mechanical states correlate with transcriptional profiles that stratify patients by clinical responsiveness and pathobiological data, uncovering a novel axis of disease associated stratification. Importantly, this work establishes that the specific mechanophenotype-associated transcriptome may serve as both a functional readout of disease state and a predictive tool for therapeutic stratification and treatment. Together, these findings highlight how the mechanomics of synovial fibroblasts, through the integration of biomechanical and transcriptomic data, can help identify specific genes directly or indirectly involved in RA pathogenesis, which may be explored in the future as potential targets for modulating cellular mechanophenotypes.

## Results

### Model of a synovial organotypic cell culture

To investigate how cellular interactions and dimensionality influence synoviocyte mechanical signatures, we established 2D and 3D mono- and co-culture systems using either healthy fibroblast-like synoviocytes (HFLS) or rheumatoid arthritis synoviocytes (RA-HFLS), alone or in combination with endothelial cells (HUVECs). 3D constructs, described in this work as organotypic cell cultures (OCCs), presented a maximum diameter of 300 µm after five days. For the co-culture, a 4:1 ratio of fibroblasts to endothelial cells was used to replicate the cell populations in the native synovium tissue, as fibroblasts have been found to be the predominant cell population in synovial RA tissue.^31^

Cells extracted from the OCCs were characterized for their mechanical phenotypes and RNA-seq was used to discriminate their corresponding mechanical gene signature in “stiff” and “soft” categories (Fig. 1A). The OCC were generated using a hanging-drop approach, yielding compact 3D cultures with reproducible size and morphology (Fig. 1B). Within five days, the OCC mature into fully developed 3D cell cultures. Light-sheet imaging revealed differences in cytoskeletal organization by staining of F-actin, where monocultures displayed more disorganized actin networks compared to their co-culture’s counterparts (Fig. 1C, Fig. S1). The maturation process can be monitored over time, and the spatial organization of synovial fibroblasts and HUVECs can be tracked through live-cell staining and imaging. Maximum intensity projections of confocal Z-stacks reveal that, by day five, HUVECs self-organize into clusters resembling early blood vessel-like structures (Fig. 1D). Given the well-documented issue of necrotic core formation in 3D cultures, primarily due to limited nutrient and waste exchange, we evaluated cell viability using live/dead staining. Our results (Fig. S2) demonstrate a minimal number of dead cells, indicating that the model maintains high viability throughout its development.

**Figure 1.**
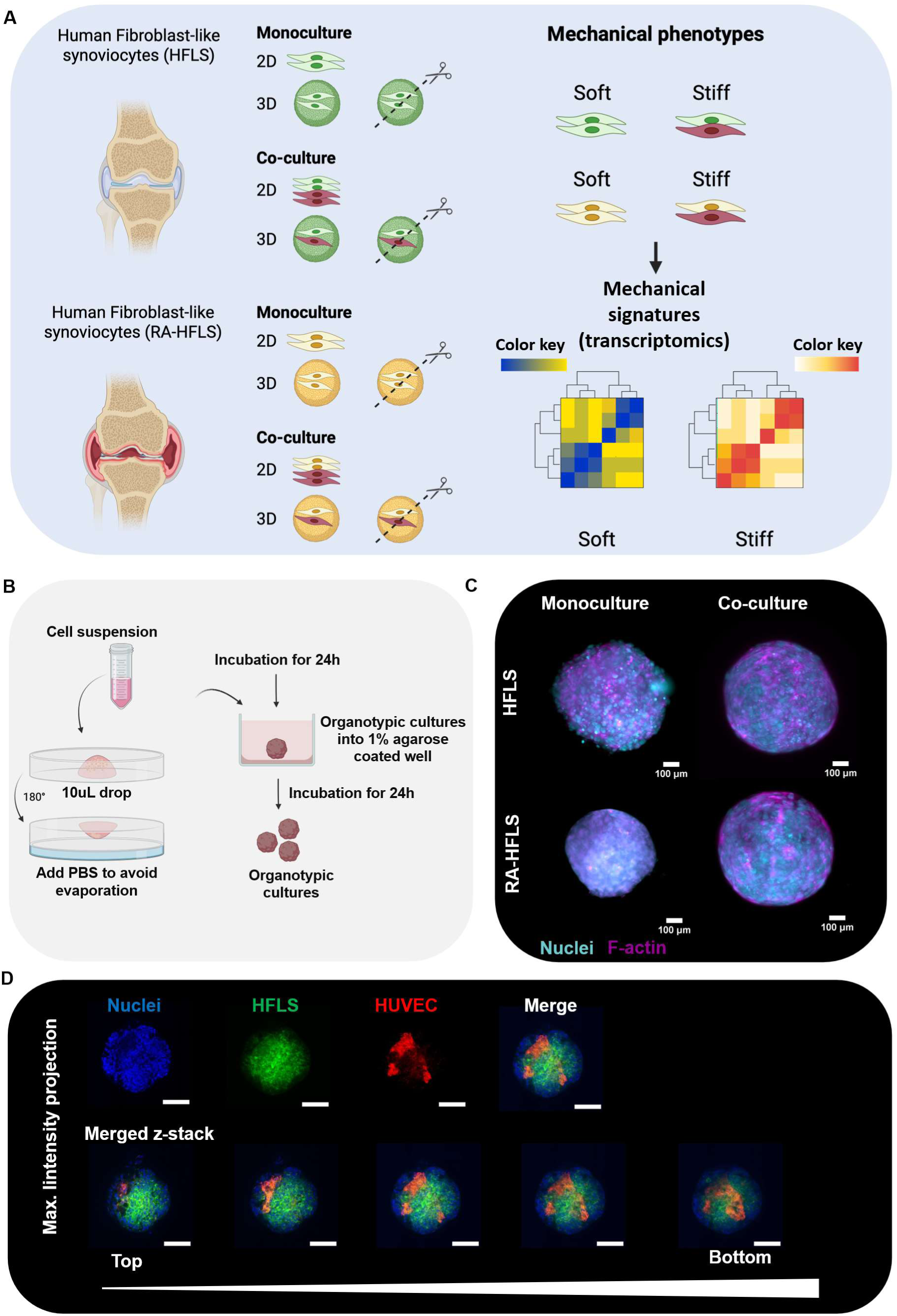
Generation of synovial organotypic cell cultures (OCC). A, B) Schematic overview of the workflow used to generate OCC and their mechanical characterizations. Scissors are representing the disassembly of the OCCs. Drops of cell suspensions were pipetted onto the lid of a culture dish and inverted to create hanging drops. After 24 h of incubation, cells aggregated to form clusters. These clusters were then transferred to agarose-coated wells supplemented with 2% basement membrane extract (BME). Within an additional 24 h, the clusters compacted to form 3D cultures. Illustration created with Biorender.com. C) Light-sheet Z-projection images of OCC stained for F-actin (purple) and nuclei (cyan), illustrating the cellular organization within the 3D cultures. D) Z-projection and Z-stack serial images of an OCC as co-culture of HUVEC (red) and HFLS (green) in a ratio of 4:1, 5 days after generation. Nuclei were stained with DAPI (blue). The z-stack images of OCC show clusters of HUVEC surrounded by HFLS. Scale bar 100 μm.

### Fibroblasts isolated from 3D organotypic cell cultures show reduced Young’s modulus

To assess one of the central biomechanical traits reflecting cell behaviour, the Young’s modulus of individual cells was evaluated under both two-dimensional and three-dimensional culture settings. The Young’s modulus, a parameter indicative of cell stiffness, was derived from high throughput nanoindentation performed on cells cultured in 2D and on individual cells isolated following dissociation of 3D organotypic cultures. Fig. 2A illustrates the distribution of Young’s modulus values for HFLS and RA-HFLS cultured as 2D and 3D monocultures. At this stage, we focused exclusively on monocultures OCC to isolate and evaluate the mechanical effects of dimensionality, deliberately excluding any mechanical contributions arising from endothelial cells. A noticeable reduction in stiffness was observed for HFLS cultured in 3D, with a median of about ∼1,3 kPa, compared to those from 2D environments with a median of ∼2,2 kPa, suggesting a softer phenotype in the 3D organotypic setting. The same experimental approach revealed that RA-HFLS also exhibited a similar trend, with significantly reduced stiffness in 3D (∼1,2 kPa) culture relative to 2D (∼1,9 kPa). However, it is interesting to notice that, RA-HFLS in general show reduced stiffness compared to HFLS from a healthy donor. Fig. 2B includes representative microscopy images taken 24 hours after transferring cells onto a 2D substrate, highlighting distinct size differences. Cells isolated from 3D cultures, both HFLS and RA-HFLS, appeared smaller compared to their counterparts from standard 2D cultures. These mechanical phenotypes can be used to profile genes associated with stiffness changes at the transcriptomic level, aiming to identify genetic signatures that regulate cell stiffness in response to the dimensionality (Fig 2C). Fig. 2D and 2E show volcano plots of up/down regulated genes in mono-culture that are in 2D or 3D cultures respectively in the healthy HFLS and RA-HFLS. However, in order to highlight the mechanical signature derived only from the dimensionality we excluded the effect of the source of fibroblast generating the volcano plot of up and downregulated genes in monocultures (Fig. 2F). Fig. 2G and 2H show the most affected genes between KEGG Pathways of differentially expressed 2D and soft 3D monoculture conditions. Genes downregulated in soft 3D culture were enriched for pathways related to TNF signaling, cytokine-cytokine receptor interaction, as well as PI3K-Akt signaling pathway. In contrast, genes upregulated in soft 3D culture were significantly enriched for pathways involved in the regulation of the actin cytoskeleton, the focal adhesions, ECM-receptor interaction, cell adhesion molecules, cell cycle and the Hippo signaling pathway. Together, these results indicate that the softer 3D culture reduces expression of soluble signaling mediators (such as TNF signaling) while promoting gene programs associated with cytoskeletal organization and adhesion-mediated mechanotransduction.

**Figure 2.**
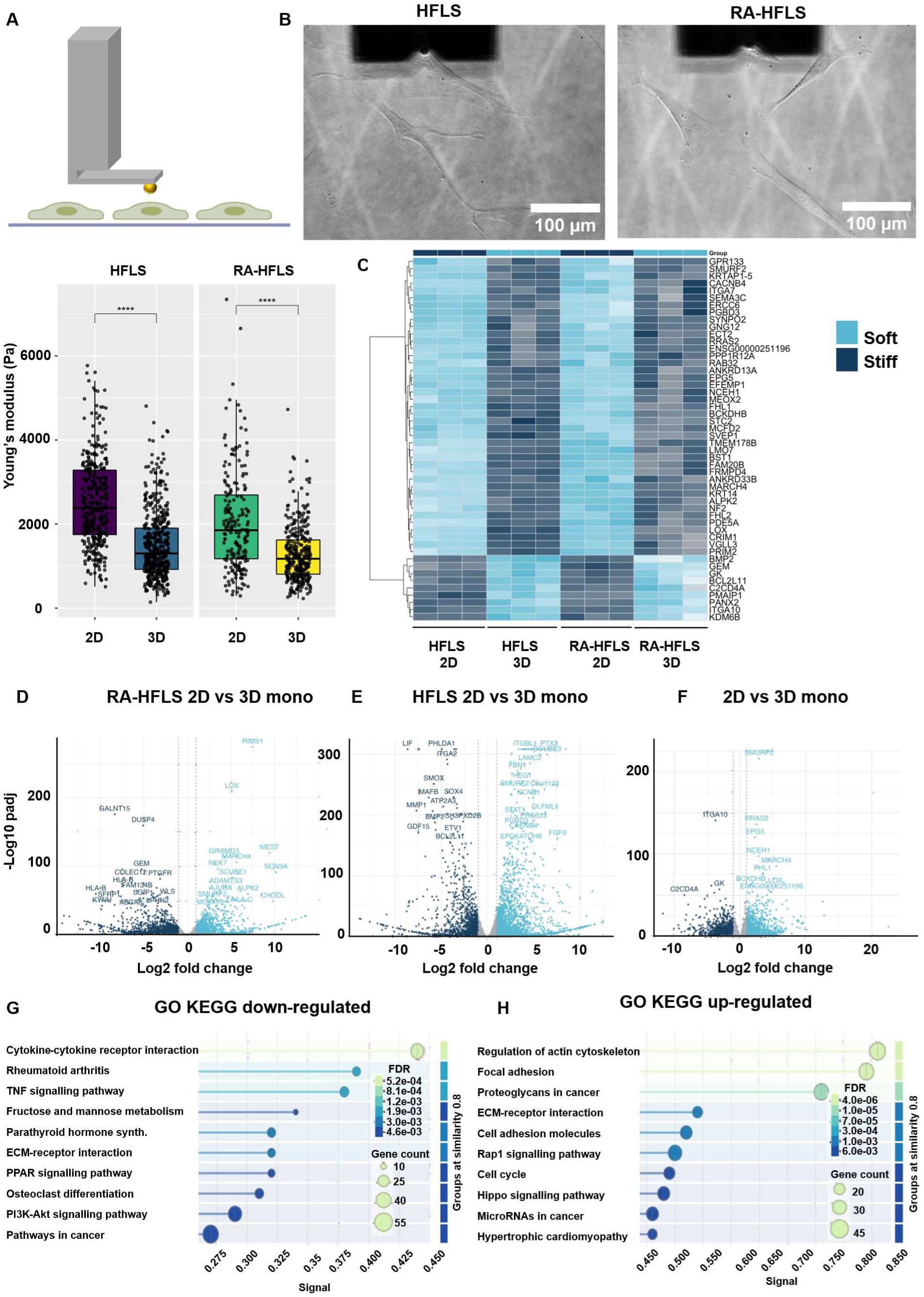
Impact of the 3D microenvironment on cellular stiffness and gene expression in OCC. A) Comparison of the Young’s modulus of single HFLS and RA-HFLS cells cultured under 2D conditions or within 3D organotypic cultures. Cellular stiffness was measured by nanoindentation. The Young’s modulus was calculated by fitting the indentation curves to the Hertz model, with >200 cells analysed per condition. Each dot represents the stiffness of an individual cell. Cells cultured in 3D exhibited significantly lower Young’s modulus values compared with cells cultured in 2D. Statistical significance between groups was assessed using one-way ANOVA followed by Tukey’s post-hoc test. B) Representative images of the nanoindentation measurement setup for cells cultured in 2D and 3D environments. Cell morphology and size differed markedly between conditions. Scale bar: 100 µm. C) Heatmap of the top 50 genes associated with soft and stiff phenotypes in HFLS and RA-HFLS cultured in 2D and 3D environments. Hierarchical clustering revealed transcriptional similarities between soft (3D cultures) and stiff (2D) phenotypes, highlighting distinct gene expression patterns linked to the different microenvironments. D, E, F) Volcano plots showing differentially expressed genes between culture conditions. Differential gene expression is shown for HFLS cultured in 2D versus 3D (D), RA-HFLS cultured in 2D versus 3D (E), and the combined comparison of 2D versus 3D monocultures independently from cell line “source” effects (F). G, H) Gene Ontology (GO) KEGG enrichment analysis of differentially expressed genes in 3D versus 2D monoculture. Downregulated genes are enriched for functions related to cytokine-cytokine receptor interaction, TNF signalling pathway, PI3K-Akt signalling pathway (G). Upregulated genes are enriched for functions associated with the regulation of actin cytoskeleton, focal adhesion, ECM-receptor interaction, cell adhesion molecules and Hippo pathways (H).

### Reduction in Single-Cell Mass Induced by 3D Culture Conditions

After observing that cells isolated from 3D cultures were smaller than those from 2D cultures (Fig. 2B), we investigated how the three-dimensional (3D) microenvironment influences cellular mass in organotypic cultures. To address this, the total mass of individual cells, cultured under both 2D and 3D conditions, were assessed using a picobalance over a 30-minute interval of time. As illustrated in Fig. 3A, HFLS maintained in 2D culture exhibited a significantly greater average cell mass compared to those derived from 3D cultures. Specifically, HFLS from 2D settings had an average mass of 5.54 ± 0.72 ng, whereas their 3D counterparts averaged 2.28 ± 1.03 ng. A similar pattern was observed for RA-HFLS: those from 2D cultures averaged 5.33 ± 1.64 ng, while those from 3D cultures averaged only 1.73 ± 0.36 ng. Thus, cells originating from the 3D organotypic cultures consistently weighed less than those from 2D environments. These observations indicate that the dimensionality of the culture environment plays a substantial role in influencing cellular mass. Fig. 3B presents time-lapse images of synoviocytes (from 2D and 3D) attached to the picobalance cantilevers, captured at the start and the end of the experiment (30 minutes) to confirm cell viability and position, which was indicated by minimal cell movements throughout the measurement period (Supplementary Video 1).

**Figure 3.**
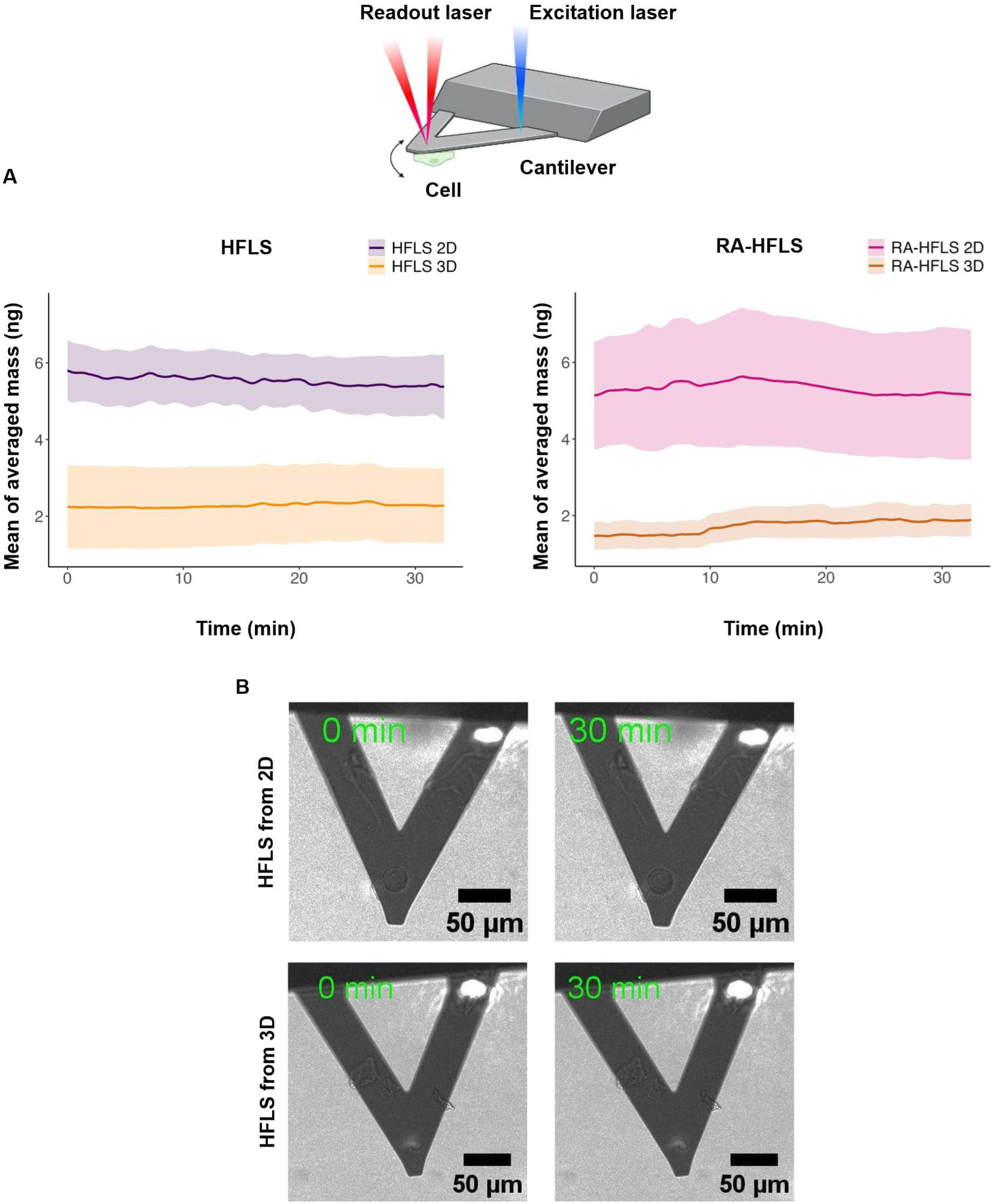
Impact of the 3D environment on total (wet + dry) cell mass. A) Comparison of the total mass of single cells derived from 2D cultures and 3D organotypic cultures. The mass of individual HFLS and RA-HFLS cells was measured using an AFM picobalance (schematic picture on top of the figure). Moving average curves were calculated for each condition. For HFLS, the average mass of cells from 2D culture (purple line) was 5.54 ± 0.72 ng (mean ± s.e.; n = 6), whereas cells from 3D culture (orange line) exhibited a reduced average mass of 2.28 ± 1.03 ng (n = 6). For RA-HFLS, the average mass of cells from 2D culture (pink line) was 5.33 ± 1.64 ng (n = 6), compared with 1.73 ± 0.36 ng for cells derived from 3D culture (brown line; n = 3). N represents the number of biologically independent cells analyzed. B) Representative time series images of HFLS derived from 2D and 3D cultures attached to a poly-L-lysine (PLL)-coated cantilever during picobalance measurements. Cells remained firmly attached without appreciable displacement throughout the measurement period, while displaying minimal movements indicating cell viability. Scale bar: 50 µm.

### Dry Mass Density of Synoviocytes in 2D vs. 3D Cultures

Quantitative phase imaging via digital holographic microscopy (DHM) was used to assess the dry mass of HFLS and RA-HFLS cultured in 2D and 3D environments. Experiments were conducted on cells obtained from disassociated OCCs aged 3 days (Fig. S4) and 5 days (Fig. 4A), with dry mass measurements taken 2 and 24 hours after cell seeding. Dry mass was normalized to cell area to calculate dry mass surface density. Two hours after seeding, both HFLS and RA-HFLS cells isolated from 3-day old 3D cultures exhibited significantly higher dry mass surface densities compared to cell isolated from 2D cultures; approximately threefold for HFLS and twofold for RA-HFLS (Fig S4). A similar pattern was observed in cells from 5-day-old 3D cultures (Fig. 4A), with mean dry mass surface density roughly 2 times higher for HFLS and 1.5 times higher for RA-HFLS relative to 2D conditions. At 24 hours post-seeding, 3D-cultured cells still showed elevated dry mass surface density, though the difference compared to 2D-cultured cells was reduced. HFLS from 3D culture maintained a 2.1-fold increase, and RA-HFLS a 1.9-fold increase. The reduction in difference over time suggests a partial adaptation of 3D-cultured cells to the 2D environment. This trend was consistent in both 3-day and 5-day organotypic cultures. Holographic reconstructions (Fig. 4B) confirmed morphological differences: cells from 2D cultures appeared larger and more spread, whereas 3D-derived cells were smaller and more rounded.

**Figure 4.**
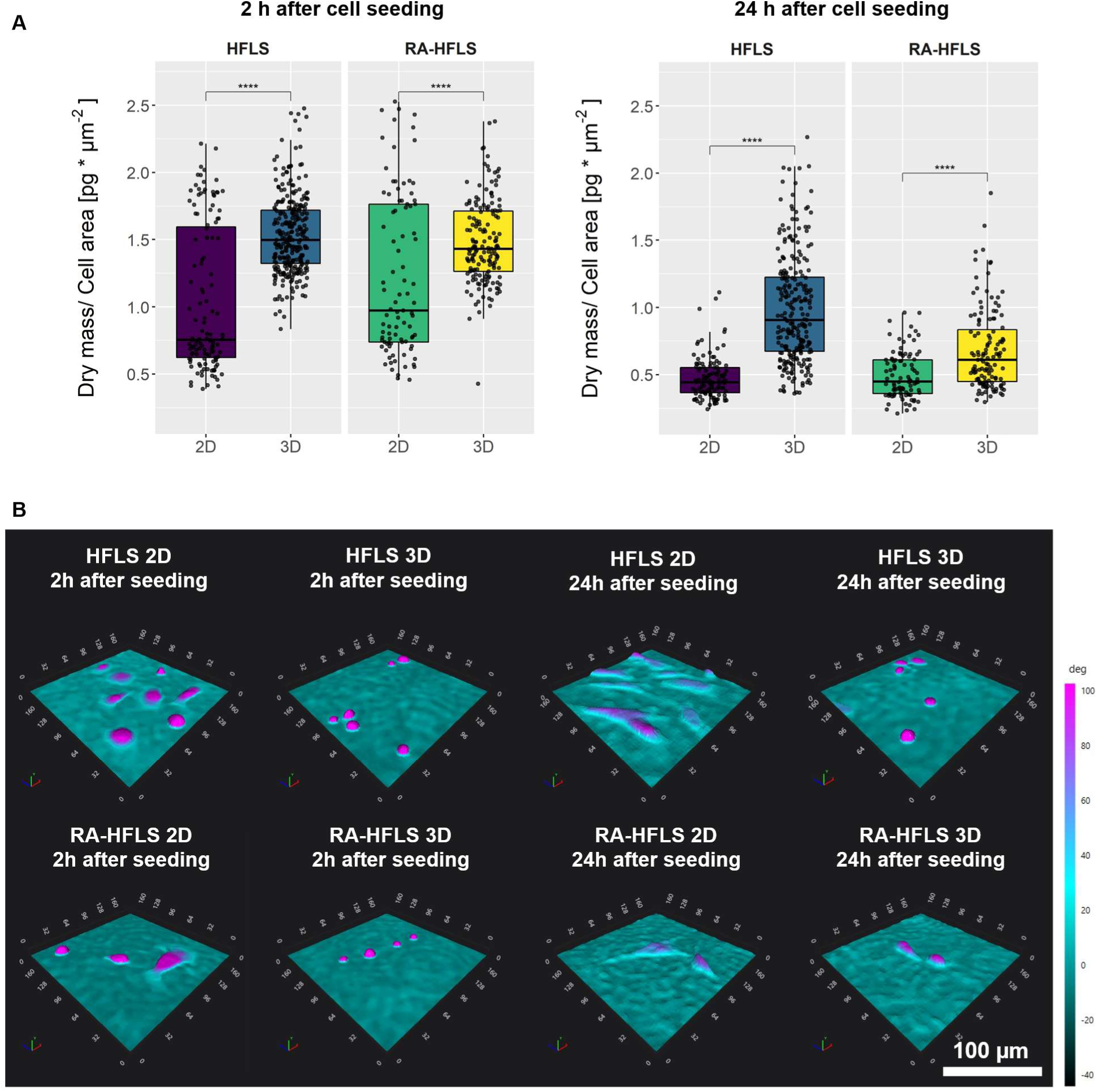
Influence of the cell environment on cell dry mass surface density. A) Dry mass surface density of synoviocytes derived from 2D cultures and 3D organotypic cultures measured 2 h and 24 h after cell seeding. The dry mass of HFLS and RA-HFLS cells was quantified using digital holographic microscopy (DHM) and normalized to the projected cell area to calculate dry mass surface density. Measurements were performed using cells isolated from 5-day-old organotypic cultures. At both time points, cells originating from organotypic cultures exhibited significantly higher dry mass density compared with cells derived from 2D cultures. Although the difference remained significant after 24 h, the magnitude of the effect was reduced compared with measurements taken 2 h after seeding. Statistical analysis was performed using one-way ANOVA followed by Tukey’s post-hoc test. B) Representative holographic microscopy images of HFLS (top row) and RA-HFLS (bottom row) derived from 2D cultures and 3D organotypic cultures. Images show cells 2 h and 24 h after seeding. Cells derived from 2D cultures appear larger and more spread, whereas cells originating from 3D cultures are smaller and display a more rounded morphology. The *n* size for each condition is between 100 and 200 cells. The statistical analysis was performed using one-way ANOVA with Tukey post-hoc test. One star (*) indicates a p-value smaller or equal to 0.05, (**) ≤ 0.01, (***) ≤ 0.001.

### Viscoelasticity of whole intact Organotypic Cultures in Mono- and Co-Culture

To evaluate the impact of endothelial cells on the mechanical properties of organotypic cell cultures, Brillouin Microscopy was used on intact 3D HFLS and RA-HFLS organotypic cultures, both in mono- and co-culture with HUVEC. The acquisition was done with a custom-made Brillouin microscope setup, briefly sketched on Fig. 5A. The setup consists of a frame incorporating a laser excitation path and a Brillouin-VIPA spectrometer. Brillouin shift, which reports on the GHz-frequency longitudinal elastic modulus of the sample, was measured using a custom-built script that fits the experimental spectra using a damped harmonic oscillator (DHO) fitting.^32^ As shown in Fig. 5B, significant differences in the normalized shift were observed between mono- and co-cultures for both cell type. The same trend can be observed in the images in Fig. 5C showing the scans of the Normalized shifts as maps. However, a clear distinction emerged also between healthy and RA-derived synoviocytes in co-culture, with RA-HFLS co-cultures exhibiting significantly higher Brillouin shift, indicating a higher longitudinal modulus phenotype associated with the pathological state. It should be noted that, while the Brillouin-derived longitudinal modulus and the Young’s modulus obtained by nanoindentation both report on the mechanical state of the sample, they probe different frequency regimes and are influenced to different degrees by hydration and compressibility. The two measures should therefore be considered complementary rather than equivalent. To further investigate this difference, staining of F-actin was performed in the organotypic cell cultures (Fig. 5D). In general, we observed that organotypic cell cultures in co-culture with HUVEC, displayed a more structured actin architecture with well-defined boundaries and internal organization (full panel of the images are shown in Fig. S1). In contrast, organotypic mono cultures showed a less defined cell arrangement, which may be linked to their lower longitudinal modulus.

**Figure 5.**
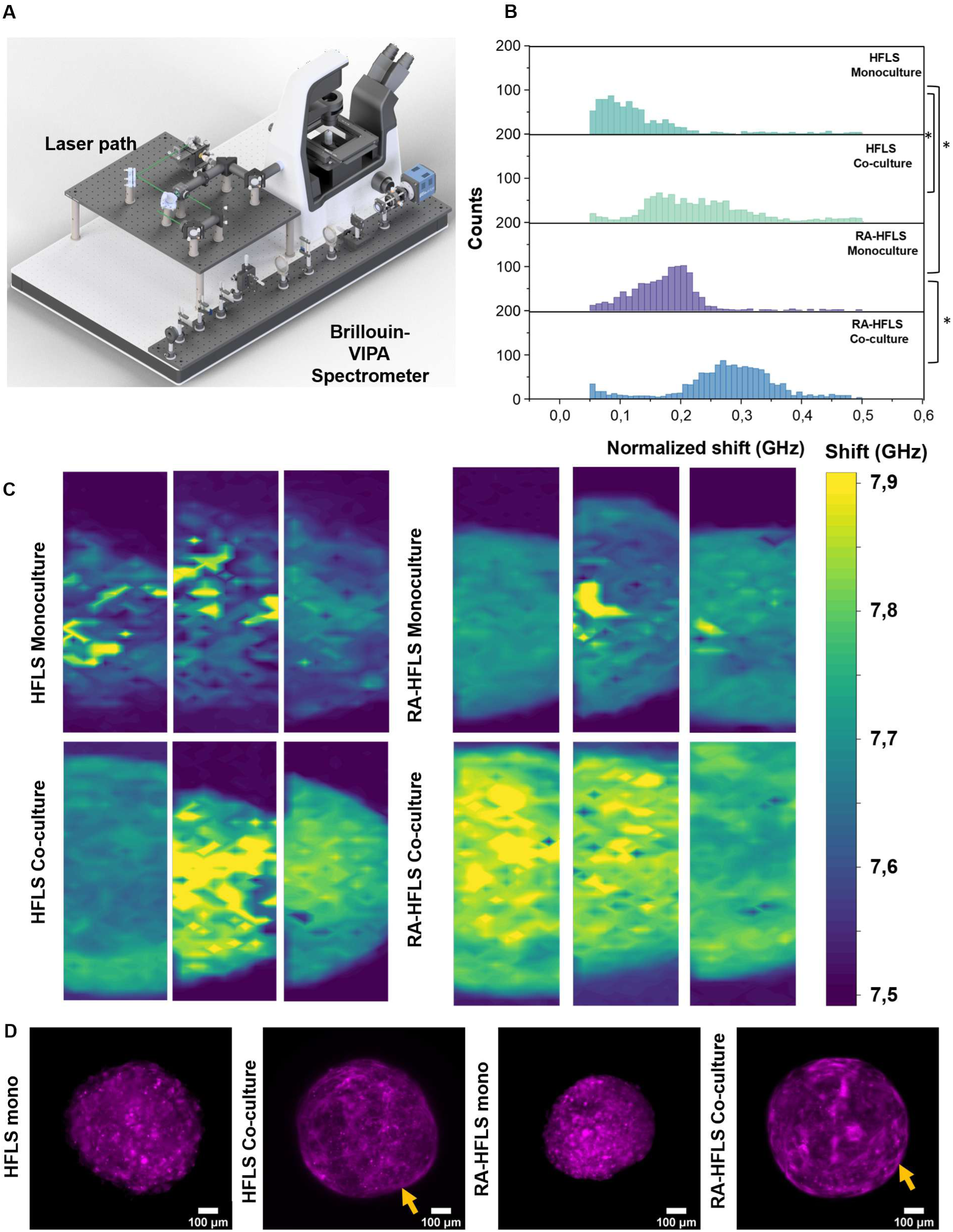
Viscoelastic characterization of whole organotypic cell cultures by Brillouin microscopy. A) Schematic diagram of the Brillouin microscope, consisting of a standard optical microscope integrated with a laser excitation path (532 nm) and a 2-VIPA spectrometer for signal detection. B) Histogram showing the distribution of normalized Brillouin shifts (GHz) across different samples; counts are reported on the y-axis. A two-way ANOVA was performed to assess statistical differences among samples, with significance set at p < 0.05. For each condition, three organotypic cell cultures were analyzed. C) Reconstruction of Brillouin shift maps for each sample, obtained over a scanning area of 50 µm x 225 µm. D) Representative image of organotypic cell cultures stained for actin. The co-culture OCCs present a better defined actin structure, as highlighted by the yellow arrows. Scale bar: 100 µm.

### Linking Biomechanical Properties and Gene Expression in Synovial Fibroblasts: Toward a Genetic Signature of the 3D extracellular environment

The aim of this work is to identify genes or genetic pathways involved in the control of the biomechanical features of synovial cells that can be used to improve patient’s stratification. To this end, we performed transcriptomic analysis for each of the conditions examined in this study and to correlate this information with a specific stiff or soft phenotype. Analysis of the Young’s modulus in monoculture conditions revealed that synovial fibroblasts isolated from 3D environments were softer compared to cells cultivated in standard flasks (see Fig. 2).

To understand the influence of Human Umbilical Vein Endothelial Cells (HUVEC) on the modulation of synoviocyte stiffness, we evaluated the Young’s modulus of HFLS and RA-HFLS under both mono- and co-culture conditions. In 2D culture, HFLS co-cultured with HUVEC exhibited reduced stiffness compared to HFLS monocultures (Fig. 6A). However, because in the co-culture both cell types were grown together in HUVEC-specific medium, we performed a control experiment to test whether the medium itself affected stiffness. HFLS maintained in HUVEC medium alone showed a comparable decrease in stiffness relative to cells cultured in standard DMEM/F12 (Fig. S3B), indicating that the reduced stiffness observed in 2D co-culture could be mainly due to the composition of medium rather than to the presence of HUVECs. In contrast, HFLS from 3D co-cultures were significantly stiffer than HFLS grown in 3D monocultures (Fig. 6A). This suggests that direct interaction with HUVEC in a 3D environment can override the softening effect of the medium and enhance cellular stiffness.

**Figure 6.**
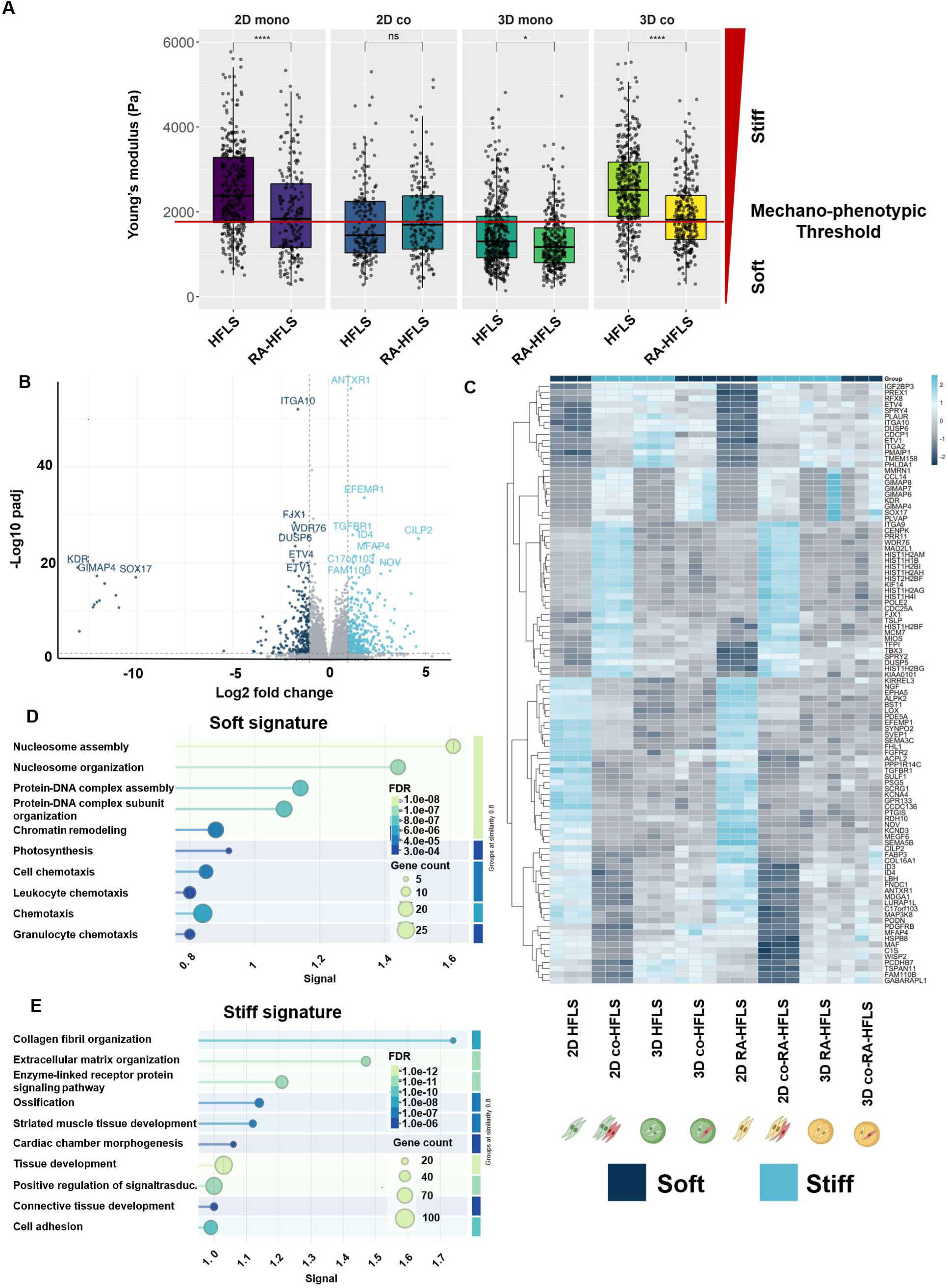
Mechanical signatures of stiff and soft fibroblast phenotypes. a) Full comprehensive box plot showing the Young’s moduli of each condition in this study. In red is shown the stiffness threshold found using statistical analysis between groups. These were used to assign stiff or soft phenotype toeach group in the RNA-seq data. B) Volcano plot showing differentially expressed genes between soft and stiff phenotypes. C) Heat map representing the up and downregulation of relevant genes across all samples and conditions. D), E) Gene Ontology (GO) KEGG enrichment analysis of differentially expressed genes in stiff versus soft phenotypes. Downregulated genes (soft phenotype) are enriched for functions related to cell cycle and DNA regulation. Upregulated genes (stiff phenotype) are enriched for functions associated with TGF-bta signaling pathway, signal pathways regulating pluripotency of stem cells, ECM-receptor interaction, Hippo and Wnt signaling pathways.

RA-HFLS displayed a different response, as no significant difference in stiffness was observed between 2D mono- and co-cultures, suggesting a lower sensitivity to medium conditions, likely due to their inflamed phenotype. However, similar to HFLS, RA-HFLS from 3D co-cultures showed a significant increase in stiffness compared to 3D monocultures (Fig. S3A, right), indicating that HUVEC co-culture in 3D affects both healthy and RA-derived fibroblasts.

To identify genes involved in the regulation of stiff and soft phenotypes independently of these factors, we pooled all Young’s modulus measurements into a single box plot (Fig. 6A), displaying all values across conditions, and defined a *stiffness-threshold* based on the statistical differences between groups. This threshold helps classify each of the conditions evaluated in this work as “stiff” or “soft”, which was then used to correlate gene expression changes. The group classification can be found on Table 1.

**Table 1.** Classification of the conditions according to mechanophenotypic threshold.

| **“Soft” phenotypes** | **“Stiff” phenotypes** |
| --- | --- |
| 2D HFSL co-culture | 2D HFSL mono-culture |
| 2D RA-HFSL co-culture | 2D RA-HFSL mono-culture |
| 3D HFSL mono-culture | 3D HFSL co-culture |
| 3D RA-HFSL mono-culture | 3D RA-HFSL co-culture |

The transcriptomic data was visualized using a comprehensive volcano plot (Fig. 6B), showing all up- and downregulated genes across samples. This plot illustrates the distribution of genes based on log₂ fold change and statistical significance (−log10 p-value). Numerous genes showed significant differential expression, with distinct clusters of upregulated (red) and downregulated (blue) genes reflecting cellular responses to culture dimensionality and presence of the endothelial cells.

Differential expression analysis revealed clearly separated gene signatures, with downregulated soft-associated genes localized on the left and upregulated stiff-associated genes on the right. Key genes associated with the soft phenotype included IL6, IL11, PTGS2, TGFB3, WNT5A, DLK1, DDIT4, HIF1A, EGR1, and BCL2L11, while the stiff phenotype was characterized by ACTA2, TAGLN, CNN1, MYL9, COL1A1, LOX, FBN1, DKK1, CTGF, MKI67, FOXM1, CCNB1, CDK1, ITGB2, TLN2, and DSP. A heatmap showing the normalised expression (z-scores) of the top 100 differentially expressed genes associated with the soft or stiff phenotypes is presented in Fig. 6C. Genes are clustered according to similarity in expression patterns across samples highlighting that the transcriptional results are highly consistent across the replicates and distinguishing stiff and soft samples.

Consistently, KEGG enrichment analysis (Fig. 6D and E) showed differential pathway activation between soft and stiff conditions, further supporting the existence of distinct mechanotransduction-related transcriptional programs. To contextualize these findings within rheumatoid arthritis (RA) pathology, the stiffness-associated gene signature was integrated with synovial tissue RNA-seq data from PEAC synovial pathotype-associated transcriptional programs.^33,34^ In the circular bar plots (Fig. 7A), genes from the soft/stiffness-associated mechanical signature were overlaid onto the clinical dataset. Genes associated with fibrotic and myeloid-fibrotic pathotypes (associated to a more aggressive pathology) are highlighted in yellow and predominantly overlap with the stiff gene cluster, whereas genes enriched in the lymphoid pathotype are shown in blue and largely align with the soft phenotype. In addition, to investigate whether mechanomic-derived gene signatures were associated with therapeutic response programs in rheumatoid arthritis genes were mapped onto the R4RA clinical trial^35,36^, a stratified phase IV randomized controlled study investigating patient responses to Rituximab and Tocilizumab in rheumatoid arthritis (RA). This analysis was performed using a three-state polar volcano plots (Fig. 7B). In this plots, angular position reflects preferential association with refractory, rituximab-responsive (Pro-RTX), or tocilizumab-responsive (Pro-TOC) transcriptional states, whereas radial distance reflects the statistical strength of association. Genes associated with the high-stiffness profile preferentially localized towards the refractory response axis (purple dots), while genes associated with the low-stiffness profile were more frequently distributed towards the Pro-RTX (red dots) and Pro-TOC response states (blue dots), or shared between these two categories (green dots). These findings suggest that biomechanical properties of synovial fibroblasts are linked to molecular programs associated with differential therapeutic response, supporting a functional relationship between tissue mechanics and clinical response in rheumatoid arthritis.

**Figure 7.**
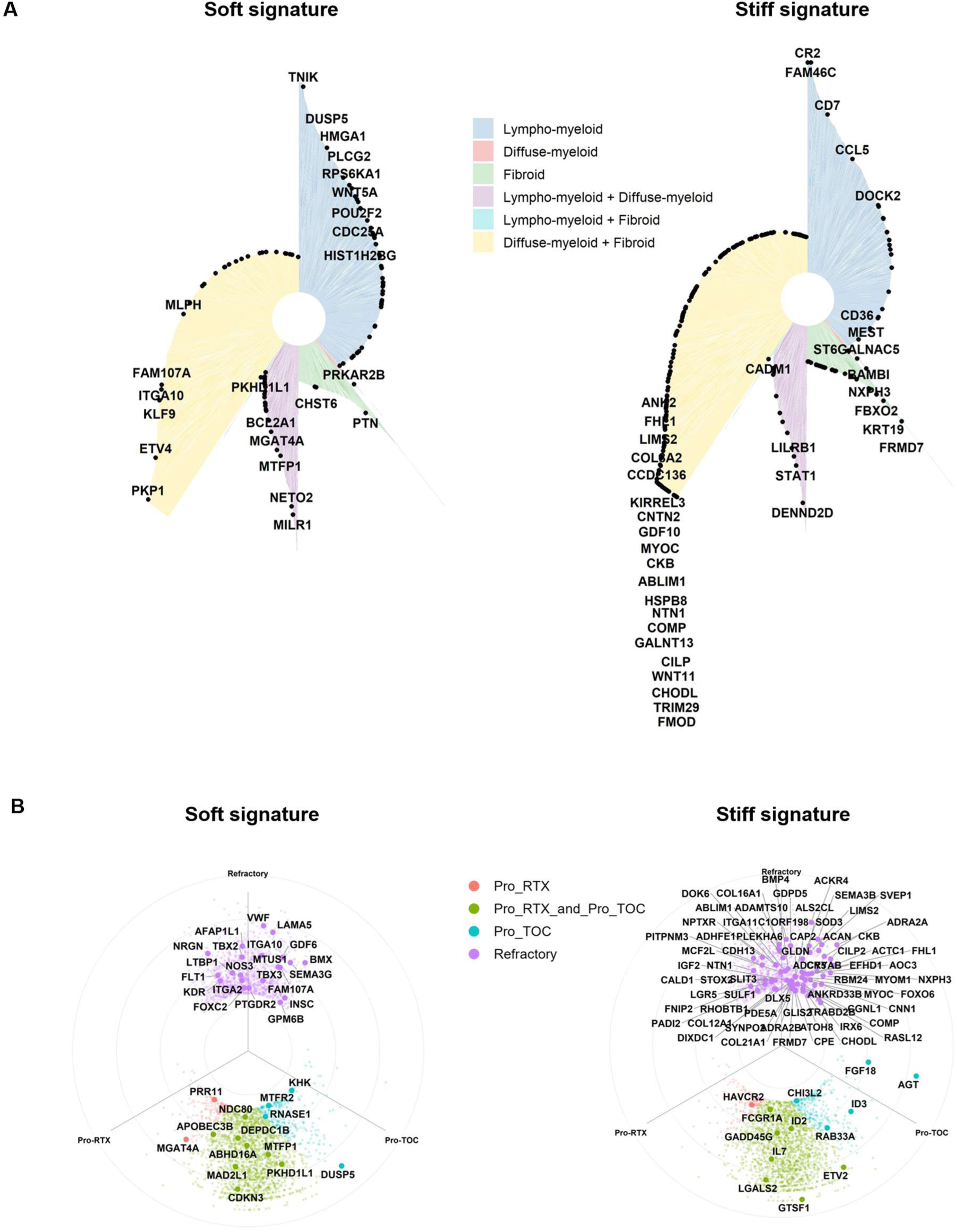
Clinical relevance of the mechanical signature in 2 public databases of RA trials. A) Mapping of the mechanical signatures onto PEAC synovial pathotype-associated transcriptional programs. Genes were assigned to lympho-myeloid (L+), diffuse-myeloid (M+), fibroid (F+) or mixed pathotype categories according to the PEAC Gene Table (see Material and Methods for details). Black dots indicate genes from mechanical signatures (stiff and soft) overlapping with PEAC pathotype-associated genes. Bar height represents the strength of pathotype association (−log10(P_LRT)). B) Mapping of the mechanical signatures onto R4RA treatment-response transcriptional programs, including refractory, rituximab-responsive, tocilizumab-responsive and shared response signatures. Colors indicate response-associated transcriptional categories and representative genes are annotated.

## Discussion

Patient stratification is fundamental for monitoring disease progression, both in tumor or autoimmune diseases. In the case of rheumatoid arthritis, a major breakthrough came with the possibility of classifying patients through biopsies characterized by their lymphocyte content (pauci-immune phenotype) or fibroblast content (fibroid phenotype). These reflect, respectively, a less aggressive or a more aggressive disease phenotype^34^. Other work has previously correlated RNA-sequencing of patients according to their response to several different drugs^35,36^. These studies combine traditional immunohistochemistry with RNA sequencing data, either single-cell or bulk, and can guide clinicians to accurately select the best drug candidates for patients, avoiding ineffective treatments.

In this exploratory work, we provide an alternative perspective from a biophysical point of view, associating an aggressive versus less aggressive phenotype based on the mechanical properties of synovial fibroblasts, the main cellular component of the synovium and key factors in rheumatoid arthritis. To our knowledge, we are the first to provide such a perspective in autoimmune diseases like arthritis. In contrast, one study has been explored in cancer, where specific mechanophenotypes have been identified and linked to altered gene expression in tumor cells. This was achieved combining machine learning-based network analysis on transcriptomic data from different tissue cell types to identify genes associated with changes in cell mechanics and validated using in silico and experimental approaches^37^. Several studies have examined transcriptional responses of specific cell types subjected to defined mechanical perturbations.^38–42^ Interestingly, comparison with these works revealed that our stiffness-associated signature includes four of the five genes identified in the mechanoregulated network reported by Urbanska et al.^37^ (CAV1, FHL2, TAGLN, and THBS1) which were likewise identified by De Marzio et al.^38^, further supporting the biological relevance and consistency of our stiffness-associated signature.

The core result was the ability to correlate a stiff and a soft phenotype with specific sets of upregulated and downregulated genes, respectively. We therefore identified specific genetic signatures that can be used for patient stratification based on biophysical information. Indeed, some genes in the stiff signature are upregulated in patients who are typically refractory to treatments such as rituximab or tocilizumab, whereas soft signature genes are found in patients who respond to these treatments. Similarly, stiff signature genes are upregulated in fibroid phenotypes, while soft signature genes are associated with lymphoid phenotypes. This opens the possibility that patients could be classified directly based on the mechanophenotype of their synovial fibroblasts, or that mechanophenotype could add an additional layer to clinical patient classification. In the future, more scalable techniques such as high-throughput nanoindentations^43–46^ and RT-DC^47,48^ could be used to assess mechanophenotypes from several hundred to million single-cells in patient-derived samples, or Brillouin microscopy could be used to perform large-scale mechanical mapping of biopsies in a label-free manner with subcellular resolution^49^.

More specifically, in this study we were able to isolate the contribution of the cellular source from other effects, focusing on the impact of the extracellular environment and dimensionality (3D environment), as shown in the related paragraph. Regardless of whether fibroblasts were derived from healthy donors or RA patients, we identified stiff and soft gene signatures independently of the cell source, but depending on 2D versus 3D conditions. This indicates a specific genetic signature linked to the culture environment: cells grown in a 3D extracellular matrix environment exhibit a softer phenotype, while those grown in 2D conditions are stiffer. Although it is well known that cells cultured on stiff substrates such as glass are stiffer than those on soft hydrogels^50^, here we associated a precise genetic signature with 3D versus 2D conditions. This is highly relevant for understanding how the 3D microenvironment can alter signaling pathways and phenotypes, making models more physiologically relevant for both developmental studies and drug testing. Among the pathways enriched in cells from 3D ECM environments are those involved in cytoskeletal regulation, focal adhesion dynamics, and receptor– ECM interactions. Previous work by Urbanska et al. showed that CAV1 is associated with stiffer phenotypes in cancer cell lines, and here we confirm this finding, as both CAV1 and CAV2 are upregulated in stiff 2D cultures (data not shown). Interestingly, in co-culture conditions, CAV1 is no longer upregulated, suggesting the endothelial cells may influence the biomechanical phenotype of fibroblasts, and will be discussed in the next paragraphs.

This study also highlights aspects related not only to stiffness but also to the regulation of total and dry cell mass. Cell mass, which is the results of the water plus the weight of the dry components such as lipids, proteins, nucleic acids, is tightly regulated and maintained in homeostasis, although it can become dysregulated under certain conditions^51^. We observed that HFLS derived from 3D environments have a total mass three times lower, suggesting that the 3D ECM acts as a constraint, maintaining smaller cell size. Similarly, cell density measured by DHM is increased in these cells after 24 hours compared to 2D cultures. Cell growth and mass regulation are known to involve the PI3K-Akt pathway, which is enriched among downregulated genes^51^. We also found the aquaporin ACQ3 upregulated in 2D, which is known as essential pathway for volume-regulatory water transport in human epithelial cells^52^, and potentially explaining differences in volume of cells. Due to technical limitations in DHM and picobalance measurements, we could not analyze co-cultures, as these methods do not allow discrimination between fibroblasts and endothelial cells simultaneously in the same sample. Furthermore, little is known about how a second cell type affects cell density regulation, making this an important area for future research.

After characterizing cells isolated from 3D ECM, we investigated the mechanical properties of entire 3D cultures. These provide global information on the combined mechanical contribution of cells and matrix. For this purpose, we used Brillouin microscopy, due to its advantages over nanoindentation and AFM. It is label-free and contact-free, allowing high-resolution mapping over large areas. It has recently been used to study biomechanical properties in spheroids,^43^ hydrogels^49^, and isomerization of small molecules.^53^ Brillouin analysis confirmed nanoindentation results on single cells extracted from different conditions, showing that the presence of endothelial cells increases stiffness in organotypic cultures with both HFLS and RA-HFLS. However, in contrast to single-cell measurements, where HFLS are generally stiffer than RA-HFLS, 3D organotypic cultures with RA-HFLS are stiffer overall. While previous studies have linked stiffness to actin organization in 3D cultures, the mechanisms by which endothelial cells are influencing the mechanical properties of the synovial fibroblasts in this context remain unclear. Based on work by K. Wei et al^54^., endothelial cells influence fibroblast positioning via Notch signaling, this suggests that these factors may contribute to the alterations in biomechanical properties following one of the previously reported mechanism known in literature^55^.

To identify a more universal biomechanical genetic signature associated with stiff versus soft phenotypes, we grouped samples solely based on nanoindentation values, using a threshold defined by statistical testing to distinguish two mechanophenotypes. The most striking result is that the stiff signature is strongly associated with aggressive arthritis phenotypes that do not respond to TOC and RTX treatments. The enriched KEGGs analysis of upregulated and downregulated gene sets reveals a shift from a proliferative phenotype, toward a mechanically active, matrix-remodeling phenotype. Specifically, the downregulated gene cohort is strongly enriched for cell cycle–associated processes, including DNA replication, mitosis, and chromosomal segregation, as evidenced by key regulators such as CDK1, CCNB1/2, CCNA2, PLK1, AURKA/B, as well as proliferation markers like MKI67, TOP2A, and PCNA. This is further supported by the downregulation of genes involved in DNA repair and replication (BRCA1, BRCA2, FANCD2, RAD51AP1), indicating a global reduction in proliferative capacity. In contrast, the upregulated gene set is dominated by pathways related to extracellular matrix (ECM) organization, cytoskeletal remodeling, and developmental signaling. Key ECM components (COL1A1, COL1A2, COL5A1, COL12A1, ELN, FBN1), matrix-modifying enzymes (LOX), and matricellular proteins (SPARC, THBS1/3, MFAP4/5) indicate enhanced matrix deposition and remodeling, processes tightly linked to increased tissue stiffness. This is paralleled by the upregulation of cytoskeletal and contractility-associated genes (ACTC1, MYL9, MYLK, CNN1, TPM1, PALLD), with increased actomyosin organization and intracellular tension. Additionally, enrichment of TGFβ, WNT, and BMP signaling components (TGFBR1, SMAD6, WNT family members, BMP4, NOG, GREM1) supports a transcriptional program associated with fibroblast activation.

## Materials and Methods

### Maintenance of Cell Lines

Human Fibroblast-Like Synoviocytes (HFLS, AcceGen Biotech ABC-TC3619) and Rheumatoid Arthritis-derived HFLS (RA-HFLS, AcceGen Biotech ABC-TC3621), sourced from single-donor, were cultured in DMEM/F-12 medium supplemented with 10% fetal bovine serum (FBS), 2 mM Glutamine, and 1% penicillin/streptomycin. Human Umbilical Vein Endothelial Cells (HUVEC, Lonza C2519A) were maintained in complete endothelial growth medium (Promocell GmbH, Germany) containing 1% penicillin-streptomycin. For passaging, cells were first washed with PBS, and subsequently incubated at 37°C in 0.05% trypsin solution for 3–4 minutes. Detachment was confirmed visually via microscopy. The enzymatic reaction was halted with complete medium, and cells were centrifuged at 300 g for 4 minutes. The supernatant was discarded, and the pellet was resuspended in fresh medium. Cells were maintained at 37°C in a humidified atmosphere with 5% CO₂.

### Generation of Organotypic Cell Cultures

A modified hanging drop method, combined with Cultrex® basement membrane extract (BME), was used to generate organotypic 3D cultures (see Fig. 1). HFLS and RA-HFLS monocultures were maintained in supplemented DMEM/F-12, whereas co-cultures (HFLS or RA-HFLS with HUVEC at a 4:1 ratio) were maintained in endothelial growth medium. To form the 3D clusters, 10–20 µL droplets containing 30,000 cells were pipetted onto the inner surface of a well plate lid. PBS was added to the wells to prevent evaporation, and the plate was incubated at 37°C with 5% CO₂ for 24 hours. Formed clusters were transferred into agarose-coated 96-well plates containing culture medium. BME was added to reach a final concentration of 2% (total volume 120 µL per well), and cultures were incubated for an additional 4 days. Medium was replaced on day 2. All experiments were conducted on day 5, unless stated differently. Brightfield images of the OCCs are presented in Fig. S5. For dissociation, organotypic cultures were washed with PBS, pooled (5–10 per sample), and incubated with 250 µL trypsin in an Eppendorf tube while shaking at 400 g for 10 minutes. Trypsin was neutralized with medium, and clusters were mechanically dissociated by pipetting. The suspension was centrifuged at 400 g for 5 minutes, and the pellet was resuspended in fresh medium for subsequent experiments.

### Light Sheet Microscopy

Scans of OCC were performed using a LS1 Live light sheet microscope (Viventis Microscopy, software v.2.0.0.3) using a Nikon ×25 NA 1.1 water immersion objective at a magnification of ×18. OCC were mounted on a single-chamber sample holder before the start of imaging. Image stacks were acquired with dual-side illumination and dual-view detection using a light-sheet thickness of 2.3 µm. Z-stacks were collected with a step size of 1 µm. Fluorescence images were acquired sequentially using 405 nm for the nuclei stained with Dapi (Sigma) and 638 nm excitation for the actin stained with SPY650-FastAct (Spirochrome), with an exposure time of 100 ms for each channel. Images were acquired at a resolution of 1024 × 1024 pixels. Image analysis was done with Fiji software.^56^ Briefly, max Z-projection were used to overlay the two different channels 405nm and 638nm. To improve image quality

### Live Cell Labeling

Live cell labeling was performed using IT-live Wheat Germ Agglutinin Alexa Fluor 594, Vybrant™ DiO, and CellMask™ Deep Red Actin Tracking Stain. Working solutions were prepared in complete media per manufacturer instructions: 1:500 for Alexa Fluor 594 and 1:1000 for Vybrant™ DiO and CellMask. Cells were incubated with the stain for 20 minutes at 37°C, followed by PBS washing.

### Viability Assay

To assess cell viability, organotypic cultures were stained with calcein AM and propidium iodide (working concentrations 1:1000 in PBS) for 15 minutes. Live cells fluoresced green, and dead cells red. Imaging was performed using an Olympus IX71 spinning disc microscope and analyzed with Fiji.

### Staining of Organotypic Cultures

Organotypic cultures were fixed with 4% PFA for 1 hour, followed by PBS washes. Blocking was performed in PBS containing 25% FBS, 0.5% BSA, and 0.5% Triton X-100 for 2 hours. Primary antibodies were diluted in PBS with 10% FBS and incubated overnight at 4°C. After sequential washes with PBS and PBS + 0.1% Tween-20, secondary antibodies were applied for 2 hours at room temperature in PBS with 10% FBS. Optional stains (DAPI, Rhodamine Phalloidin, SirActin, or Hoechst) were included as needed. Confocal images were acquired using Olympus IX71 and Nikon A1R systems.

### Measurement of Elastic Modulus via Nanoindentation

The mechanical stiffness of both cells maintained in 2D culture and cells from disassociated OCCs was assessed using the Pavone nanoindenter (Optics11 Life), after seeding in a 6-well plate and 24 hours incubation. Nanoindentation of cells was performed in the nuclear region with a 3 µm tip radius probe (0.016 N/m), using a peak load mode (max 0.01 µN; 10 µm/s speed). The cell types were discriminated by live cell labeling performed before co-culturing in 2D and OCCs. All experiments were performed at 37°C with 5% CO₂. A minimum of 200 cells were measured for each condition.

### Data Analysis Nanoindentations

Force-indentation curves were fitted with the Hertz model (Poisson’s ratio = 0.5) using DataViewer V2 software. For single cells, curves were fit to a load of 0.005 µN. Curves with R² ≤ 0.95 were excluded. One-way ANOVA followed by Tukey’s post-hoc test was used for statistical comparisons. Significance is denoted as: *p* ≤ 0.05 (*), *p* ≤ 0.01 (**), *p* ≤ 0.001 (***), *p* ≤ 0,0001 (****).

### Measurement of Total Cell Mass

Total cell mass was quantified using an inertial picobalance (Nanosurf). Cells from 2D and 3D cultures were trypsinized and seeded into petri dishes. Bruker NP-O10 B cantilevers were coated with 0.01% poly-L-lysine for 45 minutes to facilitate cell adhesion. Following calibration, each cantilever was brought into contact with a cell for 5 seconds and then lifted 300 µm to isolate the cell. Mass was recorded over 30 minutes with imaging every 5 minutes. Afterward, cantilevers were cleaned with PBS and 1% Tergazyme, followed by PBS and ethanol rinses.

### Data Analysis Picobalance

Raw data was processed in R. A moving average filter was applied to smooth fluctuations. Statistical analysis was conducted using one-way ANOVA with Tukey’s test. Significance levels were: *p* ≤ 0.05 (*), *p* ≤ 0.01 (**), *p* ≤ 0.001 (***), *p* ≤ 0,0001 (****).

### Determination of Dry Mass via Quantitative Phase Imaging

Dry mass was measured using the Lyncée Tec Digital Holographic Microscope (DHM) on a Nikon TiE inverted microscope. Around 10,000 cells from 2D and 3D cultures were seeded in glass-bottom dishes with 1.5 mL medium and allowed to adhere for 1 hour. Phase images were acquired using a 20× objective, with recordings repeated at 24 hours.

### Data Analysis Dry Mass

Holograms were analyzed with CELLS Analysis software. Background correction and segmentation were applied using a standard cell length of 25 µm. Visible artifacts were manually excluded. Data analysis was conducted only on time 0 measurements. Outliers with dry mass >1500 pg were removed. Dry mass was normalized to cell area, resulting in dry mass density (pg/µm²) to account for size variability between 2D and 3D conditions. Statistical significance was evaluated via one-way ANOVA with Tukey’s post-hoc test *p* ≤ 0.05 (*), *p* ≤ 0.01 (**), *p* ≤ 0.001 (***), *p* ≤ 0,0001 (****).

### Brillouin Microscopy

Brillouin scattering measurements were performed using a VIPA-based (virtually imaged phased array) optical Brillouin microscope equipped with a continuous-wave laser at λ=532 nm, described in detail previously by Wang et al^49,53^. A schematic representation of this microscope can be found in Fig. 5A. For this experiment, in order to scale the production and reduce size variability, the organotypic cell cultures were prepared using the AggreWell™800 24-well culture plates (STEMCELL Technologies, Vancouver, Canada). The ratio between cell populations were kept constant as described in previous paragraph “Preparation of Organotypic Cell Cultures”. Plates were prepared according to the manufacturer’s protocol. Before the measurements, organotypic cell cultures were fixed in 4% paraformaldehyde solution for 30 minutes, and stored in PBS at 4°C until analyzed. Organotypic cell cultures were placed on an Ibidi µ-Slide 8 Well high with a glass coverslip, in excess PBS, and scanned for a total area of 50 µm x 225 µm with a step size of 5 µm, encompassing a cross-section of each OCC. The exposure time of each scan was set to 800 ms. Data analysis was carried out as described previously.^57^ Statistical test was performed using two-way ANOVA followed by Tukey’s post-hoc correction (* p<0,05).

### RNA-sequencing

Five organotypic cultures per condition were collected at day 5 and centrifuged at 500 × g for 5 minutes. The supernatant was discarded, and the samples were vigorously resuspended in 50 μL of Monarch DNA/RNA Protection Reagent before being stored at −80 °C until RNA extraction. 2D samples were prepared in a similar manner: In triplicates, five samples of 30,000 cells each were seeded in a 6-well plate and cultured for five days. The culture medium was replaced after 2–3 days. After five days, cells were harvested using trypsin/EDTA, centrifuged at 500 × g for 5 minutes, and resuspended in 50 μL of Monarch DNA/RNA Protection Reagent.

Samples were then stored at −80 °C until RNA extraction. RNA was extracted using the Monarch Total RNA Miniprep Kit (New England BioLabs Inc., Ipswich, USA) according to the manufacturer’s instructions for cultured mammalian cells, including the optional DNase I treatment step. Elution was performed using 50 μL of pre-warmed nuclease-free water. The extracted RNA, without the addition of EDTA, was stored at −80 °C until sequencing. Quality control, library preparation, and RNA sequencing were carried out at the Deep Sequencing Core Facility (BioQuant, Heidelberg University). Briefly, RNA concentration and quality were assessed using a High Sensitivity D5000 ScreenTape Agilent Technology. Libraries were prepared using the Takara NGS Library Preparation Pico kit without prior RNA QC, and sequencing was performed on Illumina NextSeq 2000 platform using a P3 100-cycle flow cell. To identify differential expressed genes. The resulting FASTQ files were analyzed using RaNA-seq^58^ for preprocessing and sequence alignment, followed by downstream analysis in R-Studio using the DESeq2 package.^59^

### Synovial pathotype annotation analysis PEAC and R4RA public databases

Genes of mechanical signatures were compared with the synovial pathotype gene signatures reported by the PEAC (Pathobiology of Early Arthritis Cohort) study^33,34^. Pathotype-associated genes were retrieved from the PEAC Gene Table available through the PEAC data portal (https://peac.hpc.qmul.ac.uk/). Each gene was assigned to one of the previously defined synovial pathotype categories, including lympho-myeloid (L+), diffuse-myeloid (M+), fibroid (F+), or their combinations (L+M+, L+F+, M+F+, L+M+F+), according to the PEAC classification. Gene symbols from our dataset were matched to the PEAC Gene Table using HGNC gene identifiers. Genes present in the PEAC Gene Table were considered to be associated with the corresponding synovial pathotype group. The distribution of genes across pathotype categories was quantified by counting the number of genes assigned to each group. To visualize the relationship between the study gene set and PEAC synovial pathotypes, circular bar plots were generated in R-studio (version 2025.05.0 Build 496) using the ggplot2 package. All genes contained within the PEAC Gene Table were displayed as background bars coloured according to their assigned pathotype group. Genes from our stiff vs soft mechanical signatures that overlapped with the PEAC were highlighted as black points. For visual clarity, only the most significant genes (top 20%) within each pathotype category were annotated. Bar height was proportional to the transformed PEAC likelihood-ratio-test statistic, calculated as −log10(P_LRT), where larger values indicate stronger statistical support for pathotype association. To reduce the influence of extreme values, bar heights were capped at the 98th percentile of the −log10(P_LRT) distribution.

### R4RA treatment-response signature analysis

Genes of interest were mapped onto the synovial RNA-seq treatment-response signatures from the R4RA study^35,36^. Differentially expressed genes associated with the R4RA response groups were obtained from the supplementary R4RA gene-signature table, which classified genes into refractory, pro-rituximab, pro-tocilizumab, or shared responder signatures. Gene symbols from the study gene set were matched to the R4RA supplementary gene table using HGNC gene symbols. Genes present in the R4RA signature table were assigned to their corresponding response-associated category. The number of overlapping genes in each R4RA class was quantified. To visualize the distribution of study genes across R4RA response-associated transcriptional programs, circular and polar plots were generated in R using ggplot2. All genes from the R4RA signature table were shown as the background reference set, and genes from the study-specific list were highlighted. For circular plots, genes were assigned to a category corresponding to the PEAC study. Top 20% of the significant genes in each pathotype group are shown. For volcano-style visualizations, the x-axis represented log2 fold change for the selected comparison and the y-axis represented −log10 adjusted P value or q value. For three-way polar plots, the angular position represented the dominant response-associated direction, while radial distance represented statistical significance, calculated as −log10 likelihood-ratio-test q value. Selected genes with the strongest statistical support or biological relevance were annotated using ggrepel. This analysis was used to determine whether the study gene set preferentially overlapped with refractory, pro-rituximab, pro-tocilizumab, or shared treatment-response transcriptional programs.

## Author Contributions

**FD**: Methodology, Investigation, Validation, Formal analysis, Writing – review & editing, Visualization. **TD**: Conceptualization, Methodology, Investigation, Validation, Formal analysis, Data curation, Writing – review & editing, Visualization. **LM**: Methodology, Investigation, Data curation, Visualization. **CMG**: Methodology, Investigation, Validation. **JA, FR, RVV**: Investigation, Validation. **RF**: Formal analysis, Software, Data curation, Visualization. **FT:** Formal analysis, Software, Validation. **ZW**: Methodology, Software, Validation, Formal analysis. **CSU**: Conceptualization, Supervision, Writing, Project administration, Funding acquisition. **FC**: Conceptualization, Methodology, Data curation, Writing – original draft, Supervision, Project administration and funding acquisition. All authors reviewed and agreed to the final version of the manuscript.

## Acknowledgements

The authors would like to thank Florine Sessler and the Soft (bio)materials characterization Core Facility (Biomechanics) at IMSEAM, Heidelberg University, which is funded by the Federal Ministry of Education and Research (BMBF) and the Ministry of Science Baden-Württemberg within the framework of the Excellence Strategy of the Federal and State Governments of Germany. Funding from the European Research Council (ERC CoG No. 101001797 PHOTOMECH) is acknowledged. The authors would like to thank Moritz Rottmann and Vanessa Augustin from Leica Microsystems to help collecting images at the Viventis Microscope. F.D. and F.C acknowledges the support and funding of the Federal Ministry of Education and Research (BMBF) and the Ministry of Science Baden-Württemberg within the framework of the Excellence Strategy of the Federal and State Governments of Germany. This research has been partially funded by the Deutsche Forschungsgemeinschaft (DFG, German Research Foundation) under Germany’s Excellence Strategy via the Excellence Cluster 3D Matter Made to Order.

## Data Availability Statement

The experimental data, RNA-seq and analysis code that support the findings of this study are available from the corresponding author upon request.

## References

1. Jaalouk, D. E. et al. Mechanotransduction gone awry. Nat. Rev. Mol. Cell Biol. 10, 63–73 (2009).

2. Discher, D. E. et al. Growth factors, matrices, and forces combine and control stem cells. Science (1979). 324, 1673–1677 (2009).

3. Sun, Z. et al. Integrin-mediated mechanotransduction. Journal of Cell Biology 215, 445–456 (2016).

4. Humphrey, J. D. et al. Mechanotransduction and extracellular matrix homeostasis. Nat. Rev. Mol. Cell Biol. 15, 802–812 (2014).

5. Hynes, R. O. The extracellular matrix: Not just pretty fibrils. Science (1979). 326, 1216–1219 (2009).

6. Frantz, C. et al. The extracellular matrix at a glance. J. Cell Sci. 123, 4195–4200 (2010).

7. Engler, A. J. et al. Matrix Elasticity Directs Stem Cell Lineage Specification. Cell 126, 677–689 (2006).

8. Wei, Q. et al. Extracellular matrix in synovium development, homeostasis and arthritis disease. Int. Immunopharmacol. 121, 110453 (2023).

9. Collison, J. Extracellular matrix in the crosshairs. Nat. Rev. Rheumatol. 15, 65–65 (2019).

10. Nygaard, G. et al. Restoring synovial homeostasis in rheumatoid arthritis by targeting fibroblast-like synoviocytes. Nat. Rev. Rheumatol. 16, 316–333 (2020).

11. Bonnevie, E. D. et al. Modulating mechanobiology as a therapeutic target for synovial fibrosis to restore joint lubrication. Osteoarthritis Cartilage 32, 41–51 (2024).

12. Bartok, B. et al. Fibroblast-like synoviocytes: key effector cells in rheumatoid arthritis. Immunol. Rev. 233, 233–255 (2010).

13. Bottini, N. et al. Duality of fibroblast-like synoviocytes in RA: passive responders and imprinted aggressors. Nat. Rev. Rheumatol. 9, 24–33 (2013).

14. Neumann, E. et al. Rheumatoid arthritis progression mediated by activated synovial fibroblasts. Trends Mol. Med. 16, 458–468 (2010).

15. Yang, Q. et al. Extracellular matrix remodeling in the pathogenesis and therapeutic strategies of rheumatoid arthritis. Clin. Exp. Med. 26, 136 (2026).

16. Bartok, B. et al. Fibroblast-like synoviocytes: Key effector cells in rheumatoid arthritis. Immunol. Rev. 233, 233–255 (2010).

17. Li, H. et al. Apoptosis of Rheumatoid Arthritis Fibroblast-Like Synoviocytes: Possible Roles of Nitric Oxide and the Thioredoxin 1. Mediators Inflamm. 2013, 1–8 (2013).

18. Mousavi, M. J. et al. Transformation of fibroblast-like synoviocytes in rheumatoid arthritis; from a friend to foe. Autoimmunity Highlights 12, 3 (2021).

19. Dupont, S. et al. Role of YAP/TAZ in mechanotransduction. Nature 474, 179–183 (2011).

20. Totaro, A. et al. YAP/TAZ upstream signals and downstream responses. Nat. Cell Biol. 20, 888– 899 (2018).

21. Siebel, C. et al. Notch signaling in development, tissue homeostasis, and disease. Physiol. Rev. 97, 1235–1294 (2017).

22. Brusatin, G. et al. Biomaterials and engineered microenvironments to control YAP/TAZ-dependent cell behaviour. Nat. Mater. 17, 1063–1075 (2018).

23. Zack, S. R. et al. Another Notch in the Belt of Rheumatoid Arthritis. Arthritis & Rheumatology 76, 1475–1487 (2024).

24. Cukierman, E. et al. Taking cell-matrix adhesions to the third dimension. Science (1979). 294, 1708–1712 (2001).

25. Ravi, M. et al. 3D cell culture systems: advantages and applications. J. Cell. Physiol. 230, 16– 26 (2015).

26. Schwartz, M. A. et al. Deconstructing Dimensionality. Science (1979). 339, 402–404 (2013).

27. Colombo, F. et al. Two-Photon Laser Printing to Mechanically Stimulate Multicellular Systems in 3D. Adv. Funct. Mater. 34, (2024).

28. Chang, S. K. et al. Cartilage destruction and synovial invasion by rheumatoid arthritis fibroblast-like synoviocytes is mediated by αvβ3 integrin. Arthritis Rheum. 63, 795–806 (2011).

29. Ospelt, C. et al. Tissue destruction in rheumatoid arthritis—mechanisms, patterns and biomarkers. Nat. Rev. Rheumatol. 8, 221–229 (2010).

30. Wei, K. et al. Notch signalling drives synovial fibroblast identity and arthritis pathology. Nature 582, 259–264 (2020).

31. Lin, K. et al. Heterogeneity of active mast cells, endothelial cells, and fibroblasts in hemophilic arthritis defined by synovial single-cell sequencing. Sci. Rep. 15, 43350 (2025).

32. Bouvet, P. et al. Consensus statement on Brillouin light scattering microscopy of biological materials. Nat. Photonics 19, 681–691 (2025).

33. Humby, F. et al. Synovial cellular and molecular signatures stratify clinical response to csDMARD therapy and predict radiographic progression in early rheumatoid arthritis patients. Ann. Rheum. Dis. 78, 761–772 (2019).

34. Lewis, M. J. et al. Molecular Portraits of Early Rheumatoid Arthritis Identify Clinical and Treatment Response Phenotypes. Cell Rep. (2019).

35. Rivellese, F. et al. Rituximab versus tocilizumab in rheumatoid arthritis: synovial biopsy-based biomarker analysis of the phase 4 R4RA randomized trial. Nat. Med. 28, 1256–1268 (2022).

36. Humby, F. et al. Rituximab versus tocilizumab in anti-TNF inadequate responder patients with rheumatoid arthritis (R4RA): 16-week outcomes of a stratified, biopsy-driven, multicentre, open-label, phase 4 randomised controlled trial. The Lancet 397, 305–317 (2021).

37. Urbanska, M. et al. De novo identification of universal cell mechanics gene signatures. Elife 12, (2025).

38. De Marzio, M. et al. Genomic signatures of the unjamming transition in compressed human bronchial epithelial cells. Sci. Adv. 7, (2021).

39. Kılıç, A. et al. Mechanical forces induce an asthma gene signature in healthy airway epithelial cells. Sci. Rep. 10, 966 (2020).

40. Zhang, F. et al. Mechanomics analysis of hESCs under combined mechanical shear, stretch, and compression. Biomech. Model. Mechanobiol. 20, 205–222 (2021).

41. Strippoli, R. et al. Caveolin1 and YAP drive mechanically induced mesothelial to mesenchymal transition and fibrosis. Cell Death Dis. 11, 647 (2020).

42. Rysä, J. et al. Mechanical stretch induced transcriptomic profiles in cardiac myocytes. Sci. Rep. 8, 4733 (2018).

43. Lemahieu, G. et al. RAB5A Promotes Active Fluid Wetting by Reprogramming Breast Cancer Spheroid Mechanics. Advanced Science 12, (2025).

44. Antonelli, Y. et al. When Mechanical Stress Matters: Generation of Polyploid Giant Cancer Cells in Tumor-Like Microcapsules. Adv. Funct. Mater. 34, (2024).

45. Khan, M. et al. Dimensional memory in glioblastoma mechanics: Traction force analysis of cells cultured in 2D versus 3D collagen environments. Bioact. Mater. 55, 515–528 (2026).

46. Colombo, F. et al. Fluctuations of Dry and Total Mass of Cells Exposed to Different Molecular Weights of Polyethylene Glycol. Adv. Nanobiomed Res. 3, (2023).

47. Urbanska, M. et al. A comparison of microfluidic methods for high-throughput cell deformability measurements. Nat. Methods 17, 587–593 (2020).

48. Otto, O. et al. Real-time deformability cytometry: on-the-fly cell mechanical phenotyping. Nat. Methods 12, 199–202 (2015).

49. Wang, Z., et al. Brillouin Spectroscopy Reveals Mechanical Properties Beyond Hydration. (2026).

50. Solon, J. et al. Fibroblast Adaptation and Stiffness Matching to Soft Elastic Substrates. Biophys. J. 93, 4453–4461 (2007).

51. Fu, J. et al. Cells prioritize the regulation of cell mass density. Sci. Adv. 11, (2025).

52. Kida, H. et al. Roles of Aquaporin-3 Water Channels in Volume-Regulatory Water Flow in a Human Epithelial Cell Line. Journal of Membrane Biology 208, 55–64 (2005).

53. Wang, Z. et al. Monitoring cis-to-trans isomerization of azobenzene using Brillouin microscopy. Journal of Physics: Photonics 6, 035024 (2024).

54. Wei, K. et al. Notch signalling drives synovial fibroblast identity and arthritis pathology. Nature 582, 259–264 (2020).

55. Kasirer, S. et al. Interplay between Notch signaling and mechanical forces during developmental patterning processes. Curr. Opin. Cell Biol. 91, 102444 (2024).

56. Schindelin, J. et al. Fiji: an open-source platform for biological-image analysis. Nat. Methods 9, 676–682 (2012).

57. Zhang, J. et al. Mapping mechanical properties of biological materials via an add-on Brillouin module to confocal microscopes. Nat. Protoc. 16, 1251–1275 (2021).

58. Prieto, C. et al. RaNA-Seq: interactive RNA-Seq analysis from FASTQ files to functional analysis. Bioinformatics 36, 1955–1956 (2020).

59. Love, M. I. et al. Moderated estimation of fold change and dispersion for RNA-seq data with DESeq2. Genome Biol. 15, 550 (2014).

