## Supplementary Figures for "Mechanophenotype of synovial fibroblasts correlates with patients’ pathotypes and treatment response in Rheumatoid Arthritis"

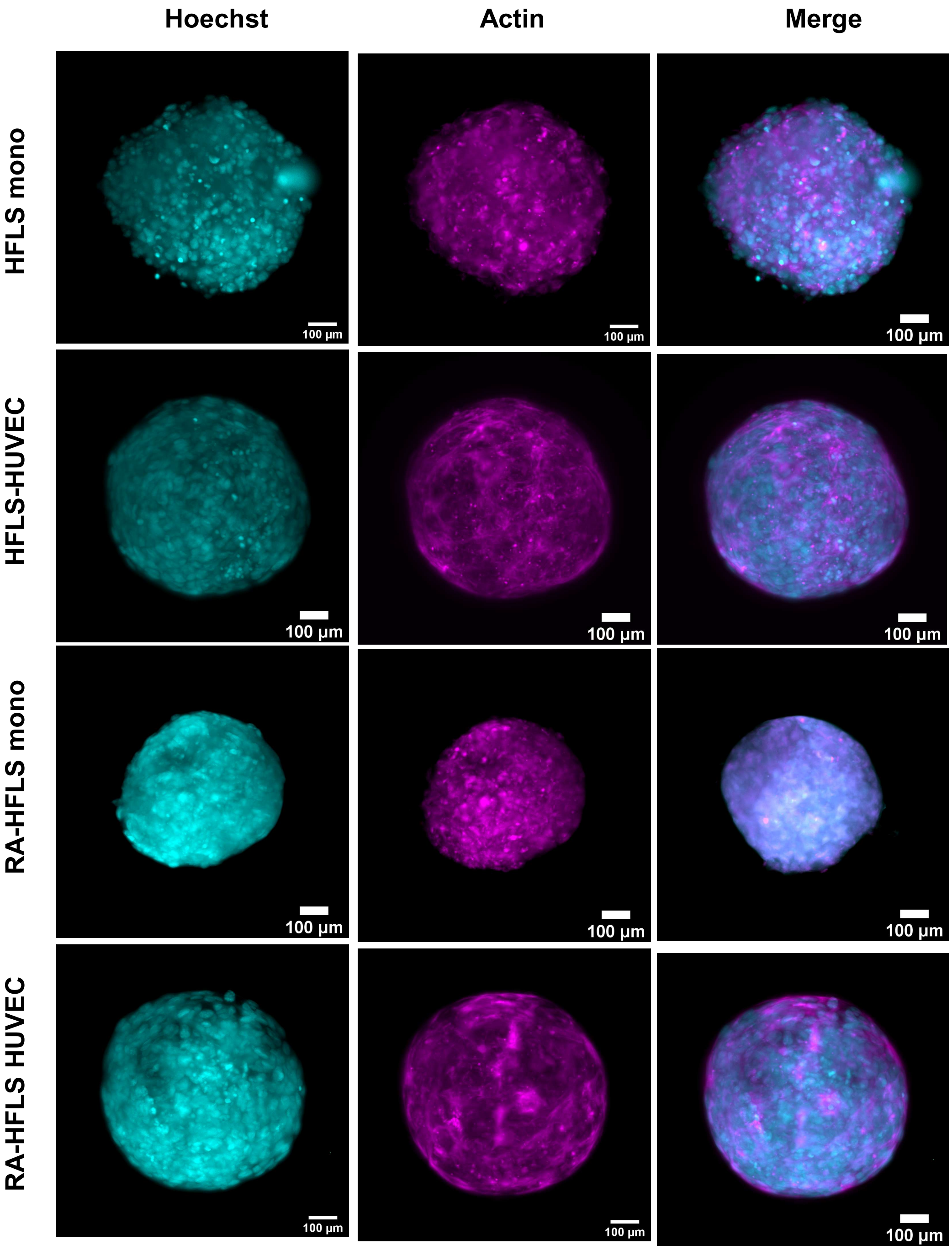


**Figure S1. Morphological differences between HFLS and RA-HFLS organotypic cultures.** Representative images of actin and Hoechst-stained HFLS and RA-HFLS organotypic cultures, in both mono- and co-culture conditions. Shown are maximum intensity projections of z-stacks with corresponding zoom-ins. Scale bars: 100 µm.

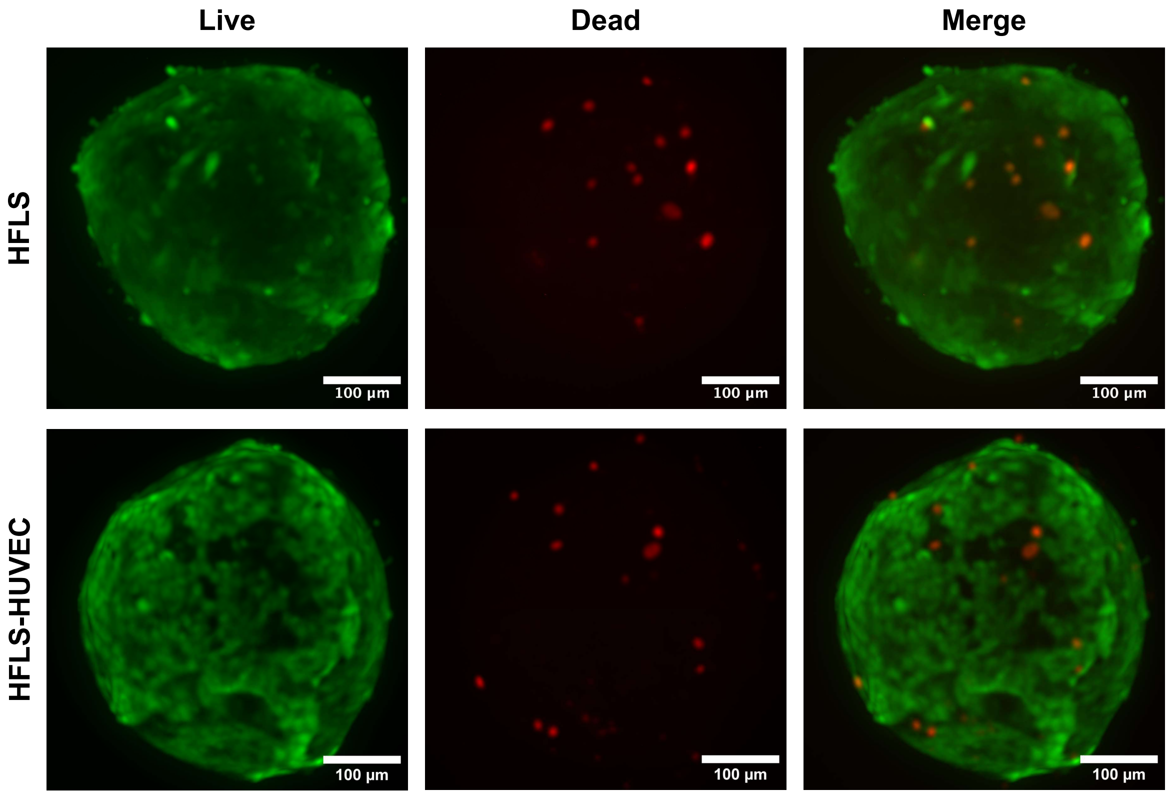


**Figure S2: Cell viability in organotypic co-cultures.** A Live/Dead assay was performed on 3D HFLS mono and HFLS co-cultures after 5 days in culture. Organotypic cultures were stained with calcein AM (green) and propidium iodide (red). For each condition 3 organotypic cultures were qualitatively analyzed and all showed a high cell viability. Maximum intensity projection of z-stack images of HFLS and HFLS-HUVEC organotypic cultures. Scale bar: 100 µm.

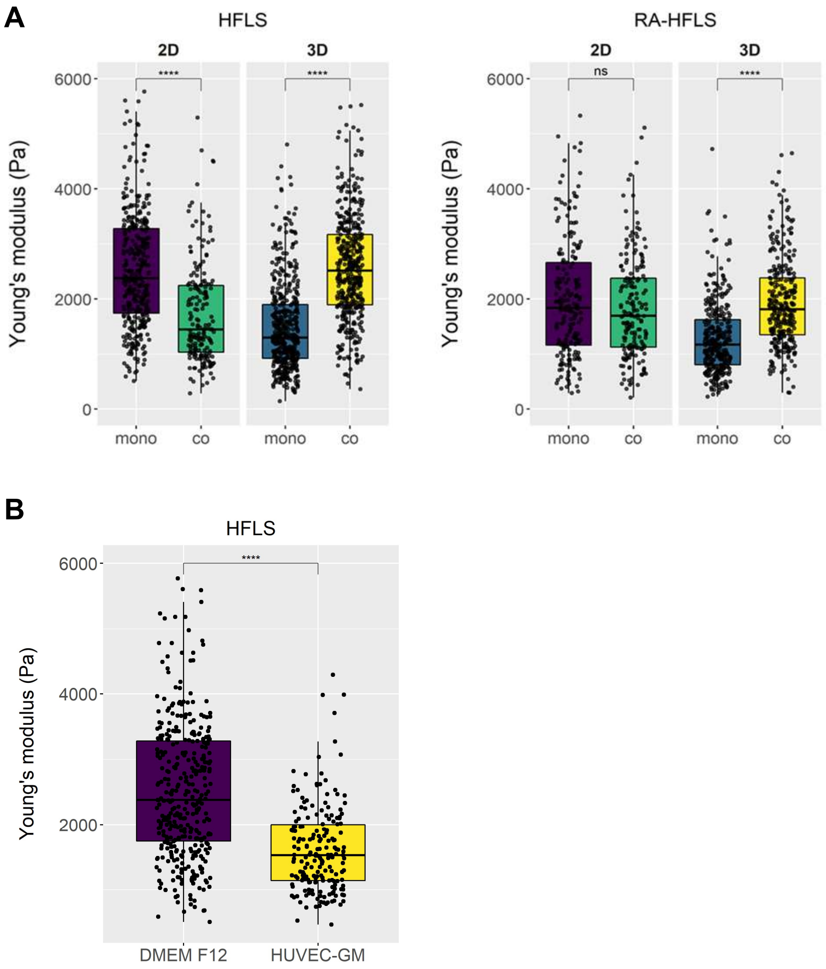


**Figure S3. Effect of growth medium on HFLS stiffness.**

Influence of Endothelial Cells and Culture Medium on Synoviocyte Stiffness. A) Young's modulus of HFLS and RA-HFLS was measured in both 2D and 3D mono- and co-culture conditions using nanoindentation. Each point represents an individual cell measurement, with >200 cells analyzed per condition. In 3D cultures, co-culture with HUVEC significantly increased cell stiffness compared to monocultures. Statistical significance was assessed using one-way ANOVA followed by Tukey's post-hoc test. The data reported in this two boxplot are based using same data plotted in Figure 6A. B) To evaluate the effect of medium composition, HFLS were cultured in either standard DMEM/F12 or HUVEC-specific medium for 5 days. Cells exposed to HUVEC medium exhibited a notable reduction in stiffness. Over 190 cells per condition were analyzed, and statistical comparisons were performed using one-way ANOVA with Tukey's post-hoc correction. HFLS were kept as 2D culture in supplemented DMEM F12 and HUVEC growth medium for 5 days. Young's modulus is significantly decreased in HFLS cultured in HUVEC medium. Each dot corresponds to the stiffness of a single cell.

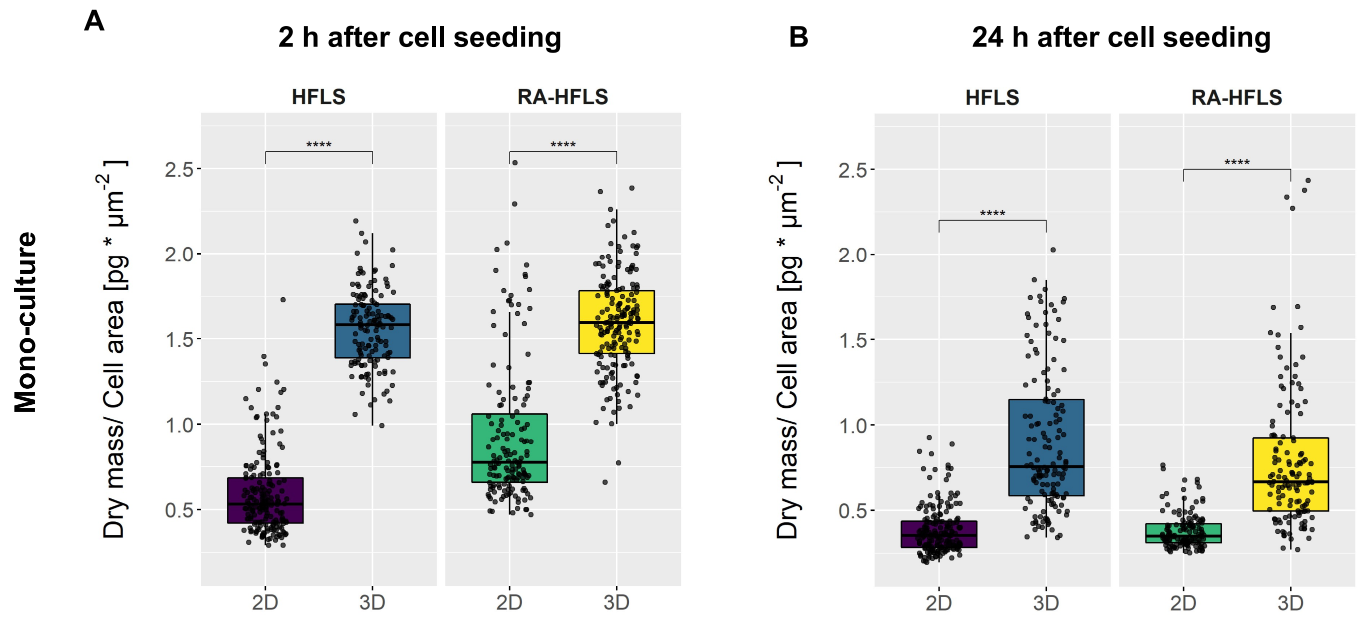


**Figure S4. Changes in dry mass / cell area ratio after 2 and 24 hours on OCCs aged 3 days.**
A), B) Measurements after 2 and 24 hours, respectively. Each dot corresponds to the measurement of a single cell. Statistical analysis was performed using one-way ANOVA followed by Tukey's post-hoc test.

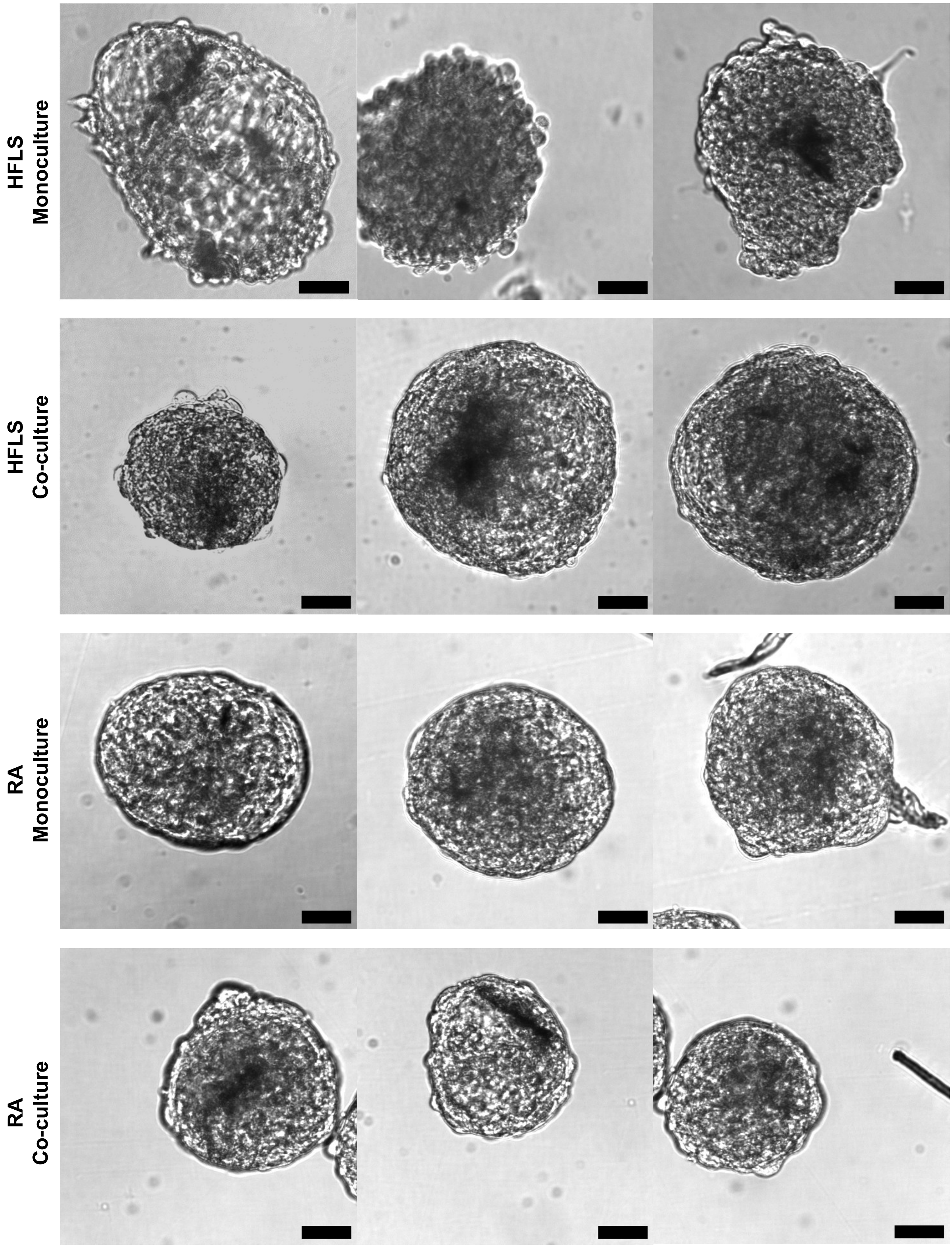


**Figure S5. Brightfield images of the organotypic cultures.**
From top to bottom: HFLS Monoculture, HFLS Co-culture, RA Monoculture and RA Co-culture, respectively. Scale bar: 100 μm
